# Knockout of *Bbs10* results in lack of cone electrical function and progressive retinal degeneration of rods and cones

**DOI:** 10.1101/2022.01.19.476952

**Authors:** Sara K. Mayer, Jacintha Thomas, Megan Helms, Aishwarya Kothapalli, Ioana Cherascu, Adisa Salesevic, Elliot Stalter, Kai Wang, Poppy Datta, Charles Searby, Seongjin Seo, Ying Hsu, Sajag Bhattarai, Val C. Sheffield, Arlene V. Drack

**Author notes:** A.T. Still University, Kirksville College of Osteopathic Medicine, Kirksville, Missouri. University of Illinois – Chicago College of Medicine, Chicago, Illinois (Megan Helms). University of Iowa Department of Biology, Iowa City, Iowa (Adisa Salesevic).

## Abstract

Bardet Biedl Syndrome (BBS) is an autosomal recessive disorder caused by mutations in at least 22 different genes. A constant feature is early onset retinal degeneration leading to blindness, with variable central obesity, polydactyly, renal failure, and developmental anomalies. BBS type 10 (BBS10) is a common form caused by mutations in the *BBS10* gene encoding a chaperonin-like protein. There are currently no treatments for the progressive vision loss. To aid in treatment development, a BBS10 mouse model was developed by knocking out the *Bbs10* gene. Using optical coherence tomography (OCT), electroretinography (ERG), and a visually guided swim assay (VGSA), we demonstrate that *Bbs10^-/-^* mice have progressive retinal degeneration. Cone electrical function was absent although cones were anatomically present on histology and retained partial function based on VGSA. The retinal outer nuclear layer (photoreceptor nuclei) progressively thinned as demonstrated on OCT and histology, and rod electrical activity decreased over time on ERG. These phenotypes are more rapidly progressive than retinal degeneration in the *Bbs1^M390R/M390R^* knock-in mouse. They are consistent with a cone-rod dystrophy distinct from typical rod-cone degeneration in retinitis pigmentosa and recapitulate aspects of retinal degeneration observed in humans with BBS10. This study has implications for BBS10 gene therapy.

## Introduction

Bardet Biedl Syndrome (BBS) is an autosomal recessive disorder caused by mutations in at least 21 different genes that are important for proper function of primary cilia (Mardy et al., 2021; Weihbrecht et al., 2017; Zhou et al., 2021). Although pleomorphic with kidney failure, obesity, developmental anomalies and diabetes, retinal degeneration occurs in all patients and may be an isolated feature (Forsythe and Beales, 2013; Green et al., 1989; Schachat and Maumenee, 1982). Retinal degeneration usually begins in the first decade of life, and results in rapidly deteriorating visual acuity, decreased visual field and eventually blindness (Green et al., 1989; Schachat and Maumenee, 1982). BBS type 10 (BBS10) is a common form caused by mutations in the *BBS10* gene encoding a chaperonin-like protein which helps to assemble the BBsome, a complex of proteins responsible for ciliary transport (Singh et al., 2020; Stoetzel et al., 2006). BBS10 represents almost 25% of all BBS cases (Forsythe and Beales, 2013), making this subtype a high yield target for treatment. Recently it has been reported that the onset and progression of retinal degeneration is earlier and faster in patients with BBS10 compared to BBS1, another common subtype (Davis et al., 2007; Grudzinska Pechhacker et al., 2021). Research into BBS10 is important to determine points during the disease course where treatment interventions would be successful.

BBS mouse models for many of the causative genes have been created to better study the etiology and potential treatment of BBS-associated retinal degeneration (Bentley-Ford et al., 2021; Cognard et al., 2015; Davis et al., 2007; Hsu et al., 2017; Zhang et al., 2011). Each BBS mouse model reported has a similar retinal phenotype to that seen in human patients (Cognard et al., 2015; Datta et al., 2015; Davis et al., 2007; Nishimura et al., 2004; Zhang et al., 2011; Zhang et al., 2013). A previous study utilized a Cre/Lox system where the LoxP sites were directed around exon 2 of the *Bbs10* gene. A total knockout was created by breeding with a Cadh16-CreDeleter line. This study explored retinal phenotype at two months of age and found thinner retinal layers and decreased phototransduction function. However, a time course of the condition was not established, differences between rod and cone function were not explored, and functional vision was not examined (Cognard et al., 2015). Extensive characterization is necessary to assess treatment effect for translational studies. Here we report the retinal degeneration phenotype of the *Bbs10^-/-^* mouse based on anatomic studies using histology, optical coherence tomography (OCT), retinal electrophysiology utilizing electroretinograms (ERG), and functional vision based on a visually guided swim assay (VGSA). The VGSA is designed to mimic the human multiluminance mobility test (MLMT) (Laird et al., 2019; Pang et al., 2008; Pang et al., 2006; Prusky et al., 2000; Sarria et al., 2015) utilized as a functional endpoint in human retinal gene therapy clinical trials. In the *Bbs10^-/-^* mouse, cone electrical activity is absent from the earliest age tested, while rod function is present though lower than normal levels and diminishes further over time. This differs from most types of retinitis pigmentosa, in which rod loss precedes cone loss. The mouse recapitulates the cone retinal phenotype reported in BBS10 patients (Grudzinska Pechhacker et al., 2021). A functional assay of cone vision, the light adapted VGSA, demonstrated that visually guided navigation is possible even with very low cone electrical function. This has implications for the possibility of recovering useful vision, even in eyes with few remaining photoreceptors.

## Results

### Deletion of the mouse *Bbs10* gene

*Bbs10^-/-^* mice were developed using an embryonic stem cell line from the KOMP repository (KOMP ES cell line Bbs10tm1(KOMP)Vlcg #052679-UCD). Bbs10 transcripts are detected in whole eyes of one month old *Bbs10^+/-^* mice, but not in eyes of *Bbs10^-/-^* mice (Fig S1). Mouse *Bbs10* is composed of two exons and both exons are deleted in this line, removing the entire protein coding region of the *Bbs10* gene. As a result, *Bbs10^-/-^* mice do not make any Bbs10 mRNA or protein (Fig. S1). *Bbs10^-/-^* males are sterile, but females are fertile, as in other BBS mice models (Cognard et al., 2015; Datta et al., 2015; Davis et al., 2007; Nishimura et al., 2004; Zhang et al., 2011; Zhang et al., 2013). Therefore, all mice must be produced with *Bbs10^+/-^/ Bbs10^+/-^* or *Bbs10^+/-^/Bbs10^-/-^* pairings using only heterozygous males and either heterozygous or knockout females. All mice were genotyped prior to experimentation and analysis. Genotyping data display a clear single 260-bp PCR product in knockout pups compared to a 445-bp PCR product in heterozygous and wildtype (WT) pups (Fig. S2). Only homozygous knockout mice display a retinal phenotype, therefore, control mice consisted of *Bbs10^+/+^* and *Bbs10^+/-^* littermates.

### *Bbs10*^-/-^ mice are smaller at birth and become progressively larger than unaffected controls

*Bbs10^-/-^* mice were smaller than WT and heterozygous *Bbs10^+/-^* littermates at 1 month of age. (Fig 1a). At age postnatal (P) day 19-21, WT pups averaged 11.11g (±1.68g Standard Deviation (SD)) (n=11) while knockout pups averaged 6.44g (±1.45g SD) (n=7). By 4 months of age the average weight of control mice was 27.43g (±4.20g SD) (n=12), while average for knockout was 36.73g (±3.35g SD) (n=3) (Fig. 1) and this difference persisted throughout life. Knockout pups had increased mortality. Special husbandry that included feeding nursing mothers and their pups Diet Gel (ClearH2O, Westbrook, ME) and delayed weaning up to P35 for survival of affected pups to adulthood. Thus, *Bbs10^-/-^* knock-out mice display weight abnormalities similar to individuals with BBS10 and required extra care to prevent mortality.

**Figure 1:**
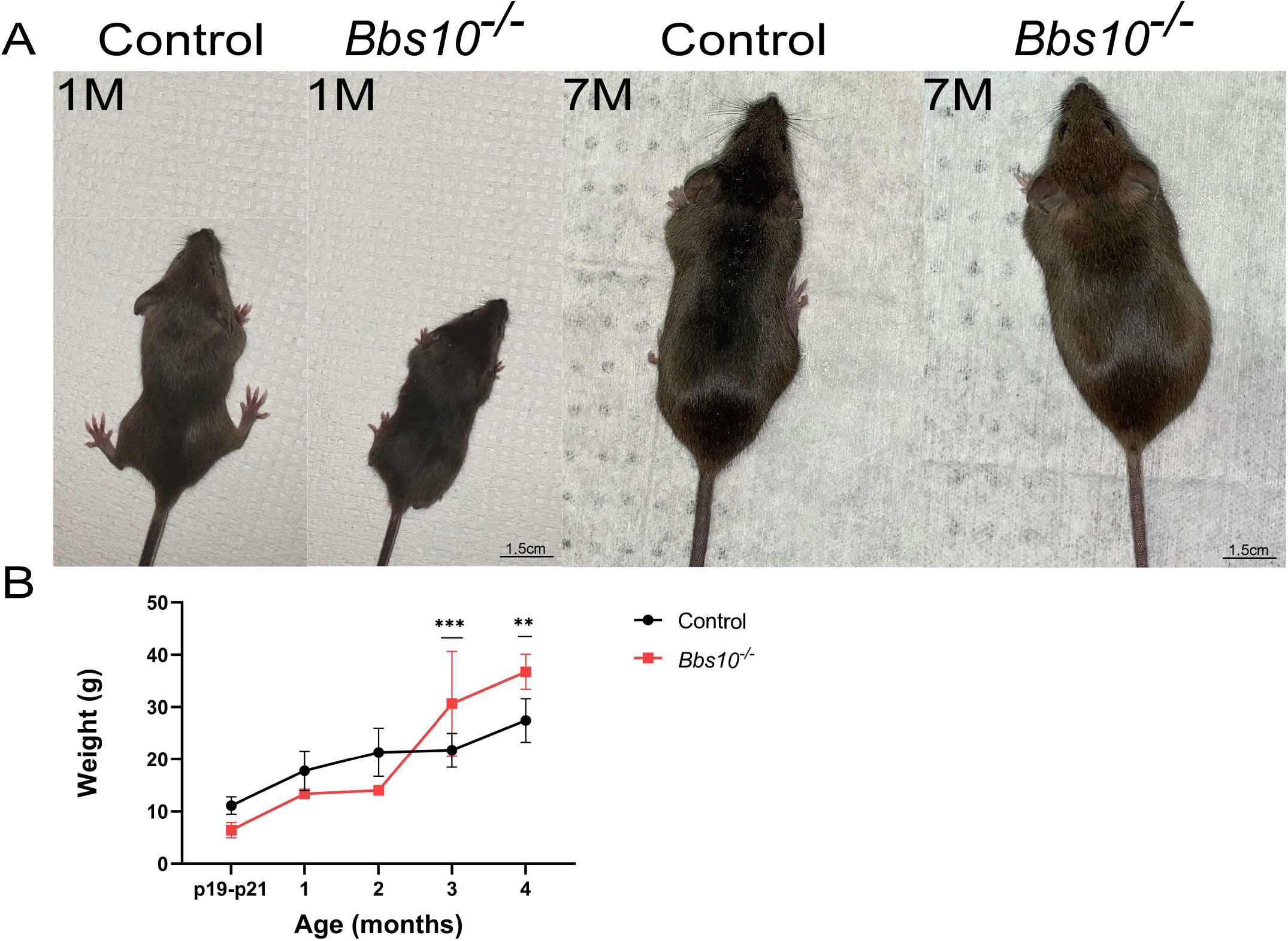
*Bbs10^-/-^* mice have altered weight relative to controls. (A) *Bbs10^-/-^* mice weigh less than control mice at birth but weigh more than control mice by three months of age. Representative mice at one and 7 months of age are shown for each genotype. Note the smaller size of the *Bb10^-/-^* mouse at one month of age and the slightly thinner abdomen of the control mouse at seven months of age. (B) Comparisons between the two mouse groups from P19 to four months of age are shown. The weights are statistically different at ages 3 and 4 months while age P19-P21 is not but this could be due to small weights and sample size. A 2way ANOVA Šídák’s multiple comparisons test was performed between the means of each age group of control and affected mice. ** = p-value<0.01, *** = p-value<0.001

### *Bbs10^-/-^* mice had progressive loss of retinal outer nuclear layer thickness

Optical Coherence Tomography (OCT) is a non-invasive imaging technique that can be used to observe the retinal layers *in vivo* (Fischer et al., 2009). The thickness of the outer nuclear layer (ONL) of the retina is a common indicator of retinal degeneration, as it contains the nuclei of rod and cone photoreceptors (Lujan et al., 2015). Loss of ONL thickness indicates a decrease in the number of photoreceptors and is associated with retinal degeneration (Fig. 2a brackets). At one month of age, the control mice had an average ONL thickness of 0.055μm (±0.002μm Standard Error of the Mean (SEM)) compared to 0.047μm (±0.001μm SEM) for *Bbs10^-/-^* mice, which is 85% of control. The thickness of the ONL for control mice remained between 0.05μm and 0.055μm from one month to 8 months of age. In contrast, the *Bbs10^-/-^* mice had progressive loss of ONL thickness each month of life. By three months of age, the average thickness of the *Bbs10^-7-^* mouse ONL was 0.028μm (±0.001μm SEM) compared to 0.054μm (±0.001μm) of control mice. This is roughly 50% thickness of that of control mice. By seven months of age, *Bbs10^-/-^* mice had no distinguishable ONL on OCT (Fig. 2b). In fact, histological analysis showed that *Bbs10^-/-^* mice retain only a single layer of cells at seven months of age (Fig. S3). Thus, OCT was able to show that *Bbs10^-/-^* mice have a significant loss of ONL over the course of their life.

**Figure 2:**
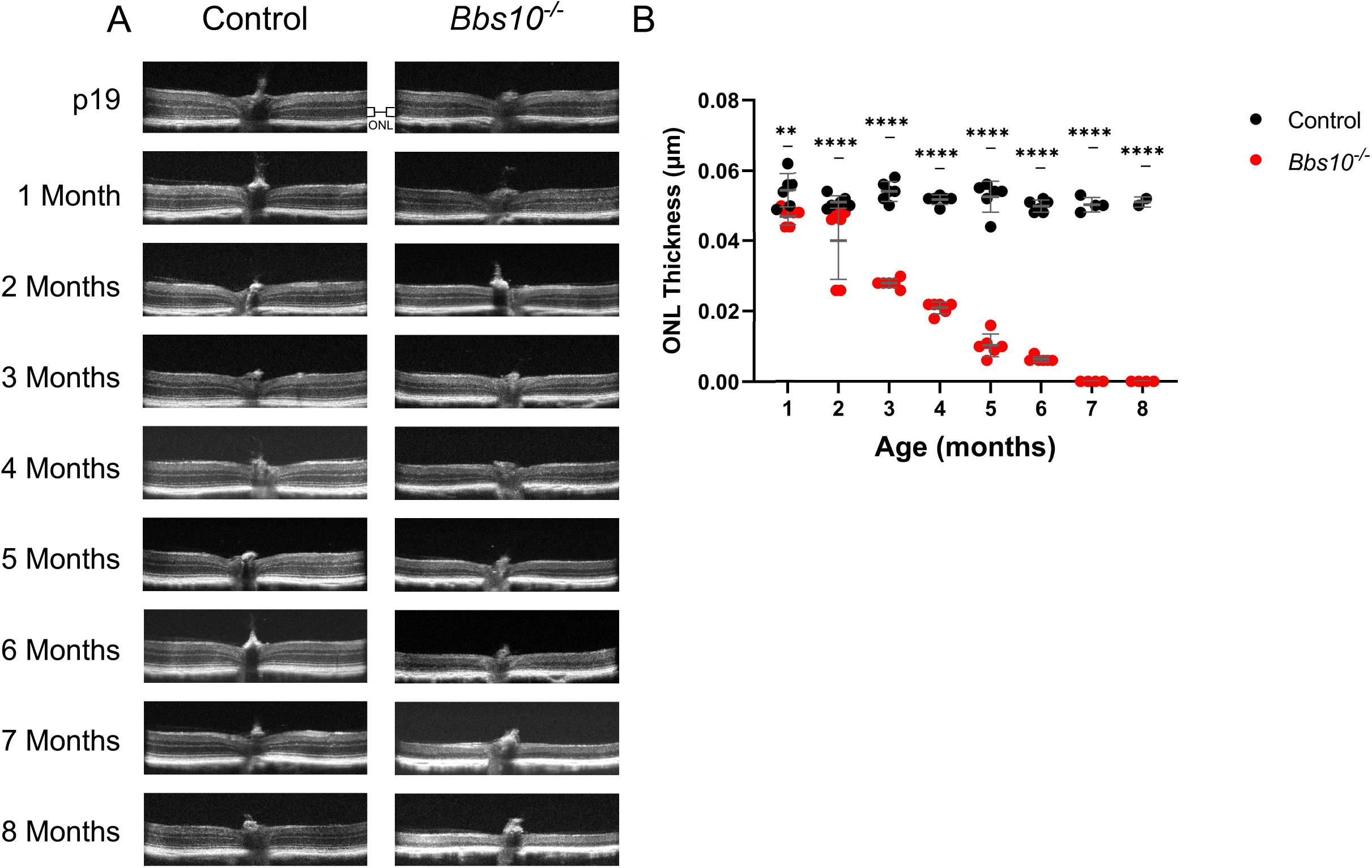
*Bbs10^-/-^* mice exhibited a progressively thinner outer nuclear layer compared to controls. (A) Optical Coherence Tomography (OCT) images of control and *Bbs10^-/-^* mice from P19 to 8 months are shown. The *Bs10^-/-^* mice exhibit a degenerating outer nuclear layer (ONL) while controls remain stable over time. (B) The graph depicts the ONL thickness of control mice and *Bbs10^-/-^* mice at different ages. The *Bbs10^-/-^* ONL was initially less than that of controls and decreased with age, until it was non-detectable at seven months of age. A 2-way ANOVA Šídák’s multiple comparisons test was performed between the means of each age group of control and affected mice. ** = p-value<0.01, **** = p-value<0.0001

### Young *Bbs10^-/-^* mice display abnormal retinal organization and lack photoreceptor cell markers

*Bbs10^-/-^* mice have abnormal photoreceptor outer segments as early as P15 shown by Transmission Electron Microscopy (TEM) (Fig. 3). By contrast, control mice exhibited organized outer segment discs of the photoreceptors. These outer segments have horizontally organized discs present in the region of the EM photo indicated by the yellow arrow. In contrast, *Bbs10^-/-^* mice lack this disc organization as well as have a lack of polarity organization in any remnant of an outer segment. This demonstrates the outer segments of the photoreceptors were anatomically abnormal in *Bbs10^-/-^* mice. This early abnormality has implications for the role of BBS10 in early development of the retina and makes early treatment of BBS10 in humans required rather than optional to target prevention and progressive degeneration of photoreceptors.

**Figure 3:**
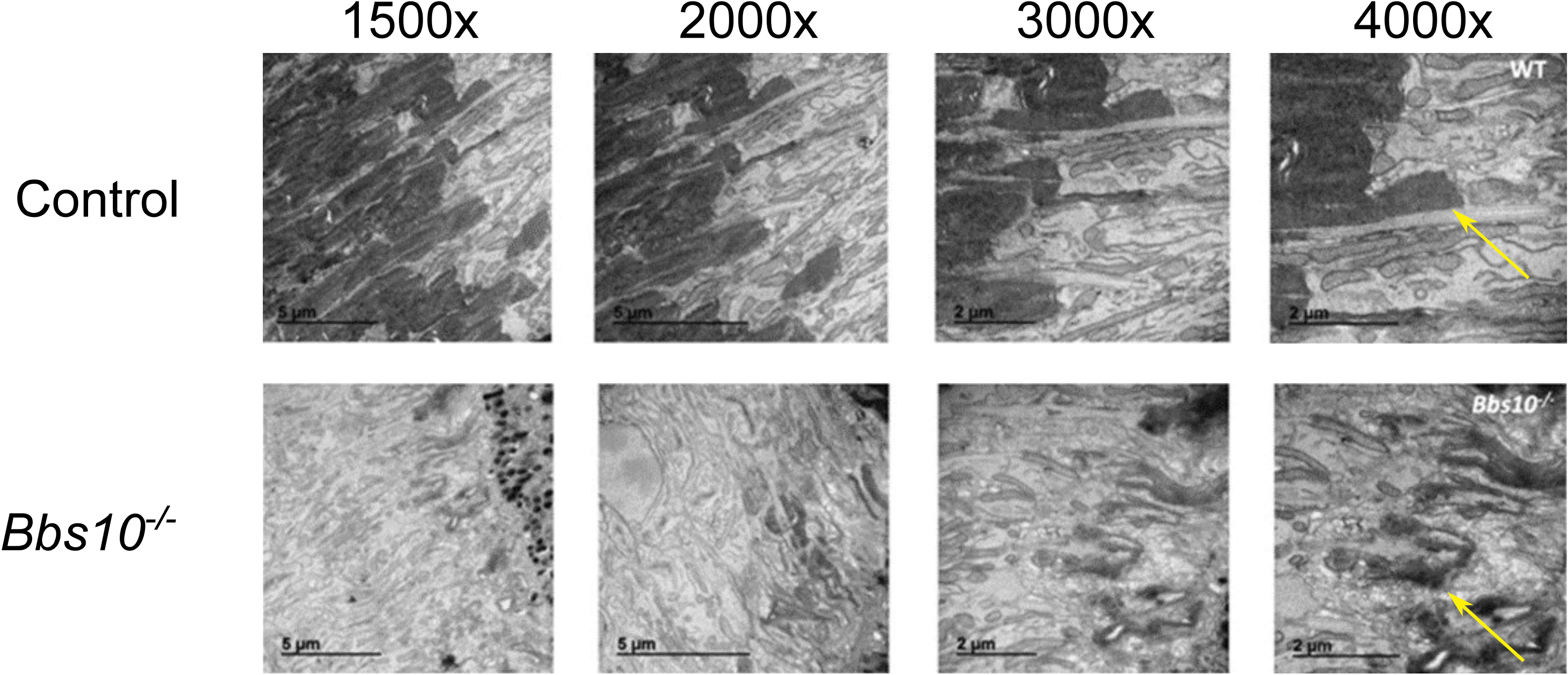
*Bbs10^-/-^* mice display disorganized photoreceptor outer segments at P15. Comparison of a control mouse retina with *Bbs10^-/-^* mouse retina under transmission electron microscopy. The large outer segment discs of the control mouse (arrow) are highly organized while the corresponding outer segments in the *Bbs10^-/-^* mouse are disorganized.

Immunofluorescence images of the retina of P21 mice reveal mislocalized and absent cone markers. OPN1MW, which is the protein pigment for Medium-wave-sensitive opsin 1, is a cone photoreceptor pigment (Deeb, 2004). It was easily detected in young P21 *Bbs10^-/-^* mice, although it was mislocalized to inner segments. This is displayed in the upper panels of figure 4. Normal localization of OPN1MW would be confined to outer segments, as seen in the control mouse retina (Datta et al., 2015). However, the cone specific protein, GNAT2, which is the alpha subunit of cone transducin (Aligianis, 2002), was almost entirely absent from *Bbs10^-/-^* mouse photoreceptor layers by P21 (Fig. 4). In previous studies of mouse BBS model, it was found that GNAT2 was highly mislocalized to the inner segment, as is the case here with OPN1MW (Datta et al., 2015). A lack of properly localized outer segments and cone specific proteins at such an early age indicate that the BBS10 disease phenotype is present very early in life.

**Figure 4:**
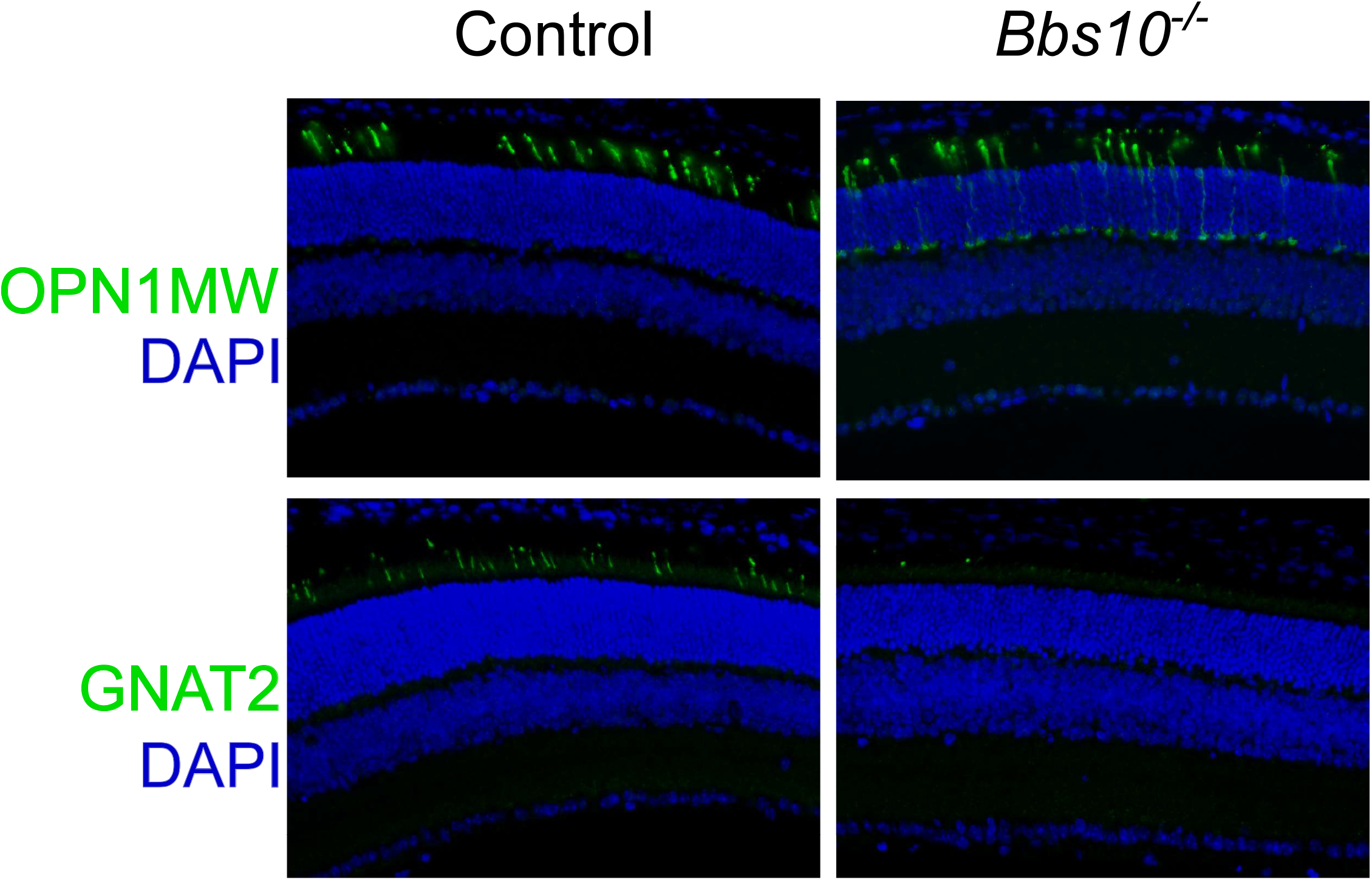
*Bbs10^-/-^* mice have abnormal cone protein localization compared to controls. OPN1MW and GNAT2 are cone specific proteins that are abnormal in *Bbs10^-/-^* retinas at P21. Control mice display prominent protein staining (green) for both cone-specific proteins. *Bbs10^-/-^* mice display staining for OPN1MW; however, it is mislocalized to the inner segments. Staining GNAT2 was nearly undetectable in mutant mice.

### Young *Bbs10^-/-^* mice display decreased electroretinogram amplitudes and lack a 5 Hz flicker response

The electroretinogram (ERG) is used to measure the entire electrical output of the photoreceptors of the retina (ref). The ERG uses a series of light flashes of different intensities in different lighting conditions to elicit separate responses from rods and cones in the retina. When dark-adapted, rods are the predominant photoreceptor that fires in response to a dim light stimulus (Robson et al., 2018). However, when light adapted, cones predominantly respond. Using a modified International Society for Clinical Electrophysiology of Vision (ISCEV) standard protocol (Mcculloch et al., 2015), four conditions were tested. Dark adapted retinas were stimulated with 0.01 cd•s/m^2^ dim flashes, to which only rods respond, followed by 3.0 cd•s/m^2^ bright flashes which elicit responses from both rods and cones (Fig. 5a and b)(Robson et al., 2018). The 3.0 cd•s/m^2^ flash is called the standard combined response (SCR) because it represents a combination of rod and cone responses. The SCR causes the largest number of photoreceptors to fire of all conditions tested and was therefore the largest response (Fig. 5b). Light-adapted conditions test cone responses and included light adapted 3.0 cd•s/m^2^ single flashes and a 5Hz flicker (Fig. 5c and d). The 5Hz flicker is used the same 3.0 cd•s/m^2^ flash, but at a high frequency of 5Hz. Cones recover faster than rods from previous light flashes (Tanimoto et al., 2015), so the 5Hz flicker elicits a response only from cones while it bleaches rods and prevents rod firing (Tanimoto et al., 2015). ERG a-waves and b-waves were analyzed for each condition; the a-waves represent primarily photoreceptor electrical firing while the b-wave is generated primarily by bipolar cells (Robson et al., 2018). Photoreceptors synapse and transmit the visual signal to the bipolar cells, thus the b-wave follows the a-wave (Robson et al., 2018).

**Figure 5:**
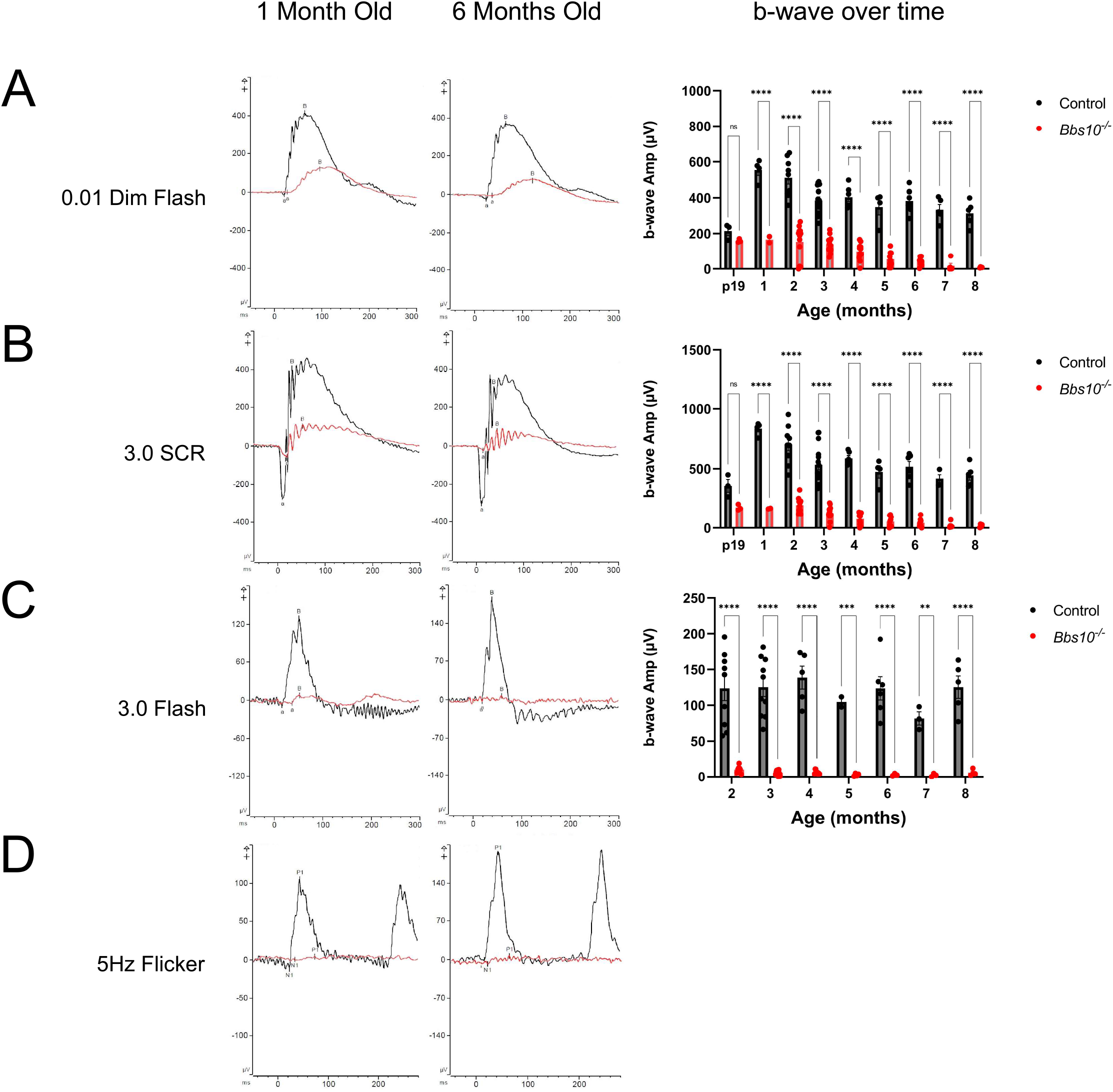
*Bbs10^-/-^* mice have abnormal electroretinograms (ERG) compared to those of control mice. (A) Overlay of a representative control and *Bbs10^-/-^* mouse ERG waveforms under the 0.01 cd•s/m^2^ dark adapted dim flash condition at one month and six months of age. *Bbs10^-/-^* mice exhibit rod generated b-waves that are recordable but consistently possess a lower amplitude than those of control mice over the course of their lifespan. (B) Overlays of representative control and *Bbs10^-/-^* mouse ERG waveforms under the 3.0 cd•s/m^2^ dark adapted standard combined response (SCR) at one and six months of age. *Bbs10^-/-^* mice exhibit b-waves with lower amplitudes when compared to those of control mice and this difference continues over the lifespan of the mouse. (C) Overlay of representative control and *Bbs10^-/-^* mouse ERG waveforms under the light adapted 3.0 cd•s/m^2^ bright flash. The *Bbs10^-/-^* mice exhibit a miniscule response at one month of age and is completely absent at six months of age. This difference is prominent throughout the lifespan of the mice. (D) The light adapted 5 Hz flicker ERG response of control and *Bbs10^-/-^* mice at one month of age and six months of age is shown. The response is unrecordable by one month of age in *Bbs10^-/-^* mice, which was the earliest time point tested. Quantification of the waveforms was not performed due to the lack of signal beyond background. A 2-way ANOVA Šídák’s multiple comparisons test was performed between the means of each age group of control and affected mice. **** = p-value<0.0001

Compared to control mice, the *Bbs10^-/-^* mice show reduced SCR ERGs by P19. At one month of age the *Bbs10^-/-^* mice are significantly different from control mice in the 0.01 dim flash and the 3.0 SCR (Fig. 5a and b). The knockout mice lack a 5Hz flicker response at this age with electrical activity of the retina indistinguishable from background signal (Fig. 5d). Interestingly, while the *Bbs10^-/-^* mice lack a recordable ERG cone response at a young age (Fig. 5c and d), cones are anatomically present on histology at P21 (Fig. 4). The *Bbs10^-/-^* mice display drastically different b-waves in the light adapted 3.0 bright flash condition. This condition became non-recordable at five months of age (Fig. 5c). The 5Hz flicker was not graphed due to the non-recordable nature at the earliest ages tested (Fig. 5d). The ERG data indicate that the *Bbs10^-/-^* mice have severe phototransduction deficiencies at an early age that become worse as they age.

### *Bbs10^-/-^* mice have decreased functional vision compared to controls

An ERG is a test of functional photoreceptors but not a test of functional vision. It is possible to have a non-recordable ERG while retaining useful functional vision. We developed a visually guided swim assay (VGSA) to test functional vision of the *Bbs10^-/-^* mouse compared to control mice. This was done by random placement of a highly visible platform in a pool of water. The mice locate this platform via vision and swim to it. Swim time was measured as time to platform (TTP), which is shorter in mice with normal vision than in mice with retinal degeneration and reduced vision.

Analysis was performed on five affected *Bbs10^-/-^* mice, and eight control unaffected mice including two unaffected *Bbs10^+/-^* mice littermates and six WT *SV129* mouse. The average TTP for control mice aged 6 months was 5.07s (±0.67s SEM) in the light and 5.50s (±0.63s SEM) in the dark. At the age of nine months, the average TTP for control mice was 3.33s (±0.39s SEM) in the light and 4.34s (±0.82s SEM) in the dark. The averages were not significantly different when compared with each other, showing that functional vision remained stable in control mice as they age. These findings also show that the visually guided swim assay provided a reproducible measure of visual function in mice.

In contrast, the *Bbs10^-/-^* mice have an average TTP at age six months of 14.08s (±3.62s SEM) while light-adapted and 20.97s (±0.68s SEM) while dark adapted (Fig 6). In both conditions, it is significantly different from WT (p-value=0.0102 and p-value=<0.0001 for light-adapted and dark-adapted respectively). It is also notable that there is a trend toward a difference (p-value=0.0982) between light and dark conditions for the *Bbs10^-/-^* mice at this age. While not significant, this suggests that *Bbs10^-/-^* mice have worse vision in the dark compared to the light at this young age. By the age of nine months, the *Bbs10^-/-^* mice average a longer TTP of 31.40s (±3.94s SEM) in the light and 35.99s (±2.57s SEM) in the dark. This is different when compared to WT of the same age and in the same lighting conditions (3.33s and 4.34s respectively) (p=<0.0001).

**Figure 6:**
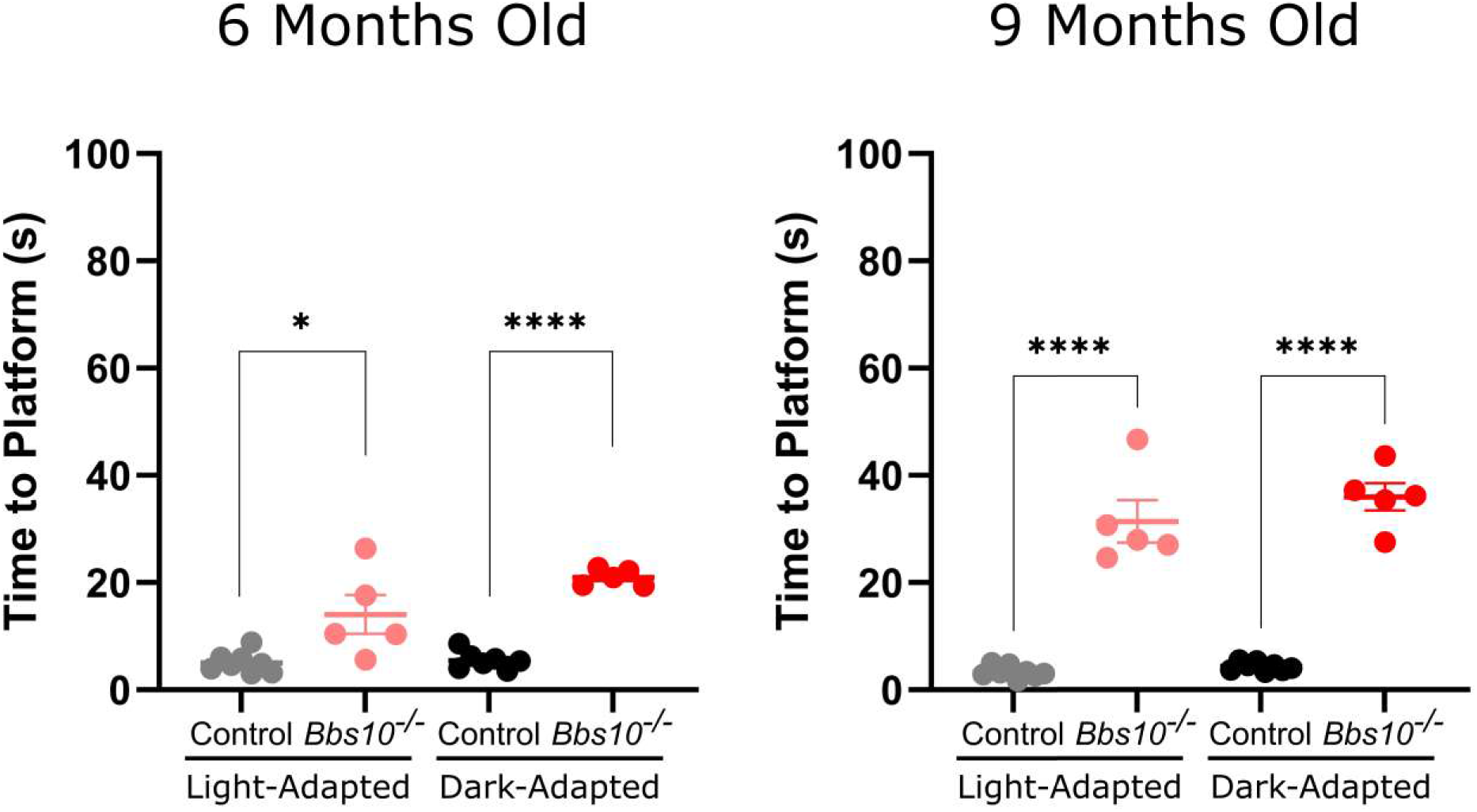
*Bbs10^-/-^* mice have reduced functional vision compared to that of control mice. The visually guided swim assay is a test of functional vision in mice and is measured by time to platform (TTP). Longer TTP suggests reduced functional vision. *Bbs10^-/-^* mice have longer TTP at six months of age in light adapted and dark adapted conditions when compared to control mice. This difference remains as the mice age to nine months. Unpaired T-tests were performed to determine statistical differences between control and *Bbs10^-/-^* mice at different ages and light conditions. * = p-value<0.05, **** = p-value<0.0001

The *Bbs10^-/-^* mice also display progressive vision loss. The difference in TTP between young and old *Bbs10^-/-^* mice was significant for the light-adapted and dark-adapted conditions (p=0.0119 and p=0.0005, respectively). The VGSA is more sensitive than ERG in detecting cone function. The initial ERG of the 5 Hz flicker showed that the cone response was already below the threshold of detection of the full field ERG, which is a mass response. This indicates that a small cohort of functioning cones were not detectable on ERG because their summed amplitude was not robust enough for detection, however this same small cohort of functional cones was adequate for essentially normal navigation in the swim assay in young mice. As the mice aged and the number of cones declined, the VGSA was able to detect a loss of function due to fewer photoreceptors, even though ERG waveform amplitudes were already indistinguishable from background signals.

### *Bbs10^-/-^* mice exhibit a more severe retinal degeneration than the *Bbs1^M390R/M390R^* BBS1 mouse model

In humans, Bardet-Biedl Syndrome Type 1 (BBS1) has been shown to have a less severe disease progression than BBS10 (Grudzinska Pechhacker et al., 2021). Together, BBS1 and BBS10 account for about 50% of all BBS cases (Grudzinska Pechhacker et al., 2021). A mouse model of the most common BBS1-causing mutation, homozygous M390R, was developed by our group previously (Davis et al., 2007) and was available for comparison to the BBS10 mouse model described here. For this comparison, both mouse models were crossed onto the 129/SvJ background for at least 7 generations. At six months of age, the *Bbs10^-/-^* mice were compared to *Bbs1^M390R/M390R^* mice. *Bbs10^-/-^* mice completely lack the ONL at six months of age. By contrast the *Bbs1^M390R/M39tlR^* retain a thin but present layer on OCT images at six months of age. ERGs taken from six-month-old *Bbs1^M390R/M39tlR^* mice show distinctly higher waveforms compared to waveforms from *Bbs10^-/-^* mice. Of note, a robust 5Hz flicker response is present in young *Bbs1^M390R/M390R^* mice and remain detectable at six months, while the 5 Hz flicker response is absent from *Bbs10^-/-^* mice even at a young age (Fig. 5d and Fig. 7b). Functional vision of *Bbs1^M390R/M390R^* mice is also better at six months of age than age-matched *Bbs10^-/-^* mice. In light adapted and dark adapted conditions, results demonstrate that the mouse models recapitulate the humans diseases, with BBS10 being more severe than BBS1.

**Figure 7:**
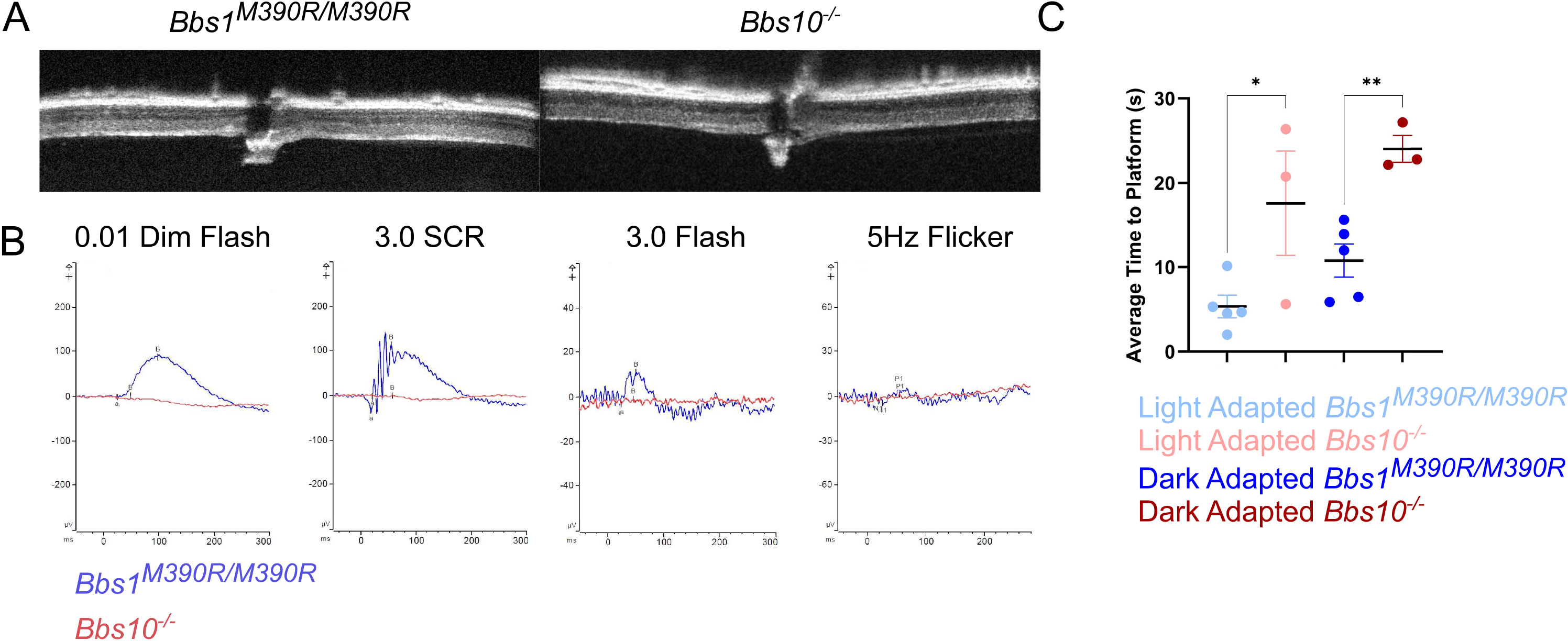
*Bbs10^-/-^* mice have a more severe retinal degeneration phenotype than that of a BBS1 mouse model, *Bbs1^M390R/M390R^*. (A) OCT images of six months old *Bbs1^M390R/M390R^* mice and *Bbs10^-/-^* mice show differences in ONL thickness. The *Bbs1^M390R/M390R^* mouse had a thin but present ONL while the *Bbs10^-/-^* mouse lacked any observable ONL. (B) ERG waveform overlays of the four ERG conditions previously described (Fig. 5) between *Bbs1^M390RM39R^* and *Bbs10^-/-^* at six months of age are shown. *Bbs10^-/-^* mice have lower waveforms than *Bbs1^M390R/M390R^* mice in all four conditions. *Bbs1^M390R/M390R^* mice have a detectable 5 Hz flicker response at six months while the *Bbs10^-/-^* lack this response from at least one month of age onward. (C) *Bbs1^M390R/M390R^* mice exhibit better functional vision compared to that of *Bbs10^-/-^* mice at six months of age in the VGSA. The TTP for *Bbs10^-/-^* mice in both light adapted and dark adapted vision is longer than that of the *Bbs1^M390R/M390R^* mice tested under the same conditions. Unpaired T-tests were performed to determine statistical differences between *Bbs1^M390R7M390^* and *Bbs10^-/-^* mice at different ages and light conditions. * = p-value<0.05, ** = p-value<0.01

## Discussion

This study sought to characterize the retinal phenotype in a new mouse model of BBS10, one of the most common types of BBS in humans (Grudzinska Pechhacker et al., 2021; Stoetzel et al., 2006). The mice display a characteristic progressive retinal degeneration that is anatomically and functionally distinguishable from unaffected control mice, as well as from a BBS1 mouse model.

The lack of response in the cone ERG could be a result of loss of functional cone transducin, needed for transmitting signals from cone photoreceptors (Aligianis, 2002), or the amplitude of the electrical signal may be lower than the threshold of detection on ERG. This could be due to either low numbers of cones firing or low amplitude of the electrical impulse from individual cones. Our discovery of disordered outer segments and lack of GNAT2 protein at early ages when the ERG is still recordable in *Bbs10^-/-^* mice suggest an early developmental need for Bbs10 protein in photoreceptor formation as well as maintenance. It is known that levels of BBS proteins start to increase around P4-6 and peak at P15 when the outer segment begins to mature (Hsu et al., 2017). The levels of proteins begin to decrease after P15 to maintenance levels (Hsu et al., 2017). The abnormal anatomical phenotype in the *Bbs10^-/-^* mice suggests Bbs10 is also important during this early development period. It can be speculated that the reason for the mislocalization of OPN1MW and complete lack of GNAT2 is a result of BBSome malfunction at P15. It is possible that at P15, OPN1MW needs the functional BBSome for transport into the outer segment. GNAT2 is a protein that forms a complex with two other subunits to produce a full functional protein, cone transducin (Aligianis, 2002). It may need BBS10 to fold properly and bind with its partners. Without BBS10 to direct formation of a functional BBSome, cone transducin cannot form in these early stages of eye development and is therefore completely absent at in photoreceptors at P21.

*Bbs10^-/-^* mice perform the VGSA at six months with faster TTP times in bright light than in dark light, averaging much closer TTP to control mice in the light than in the dark. This indicates that functional vision in bright light, remains present early in life, despite the lack of GNAT2 in cones and lack of a 5Hz cone response on ERG. Because the ERG is a mass retinal response and there are far more rods than cones in the mouse retina (Fu and Yau, 2007), a reduction in rod numbers may result in a diminished ERG amplitude, while a proportional reduction in cone numbers may result in an electrical output too small for our current technology to record. Our data suggests that very few functional cones are needed to enable useful vision in the light. This is promising for treatments such as gene therapy because it suggests that reactivating or rescuing even a small number of cones in humans with BBS10 would confer a large functional benefit.

Although mutations in BBS disease genes generally cause syndromic retinal degeneration, isolated retinitis pigmentosa (RP) has been reported with mutations in *BBS1* (Estrada-Cuzcano et al., 2012a; Estrada-Cuzcano et al., 2012b) and *BBS10* (Grudzinska Pechhacker et al., 2021). The retinal phenotype of BBS1 is typically described as retinitis pigmentosa-like rod-cone dystrophy (Grudzinska Pechhacker et al., 2021); however, BBS10 may occur as a cone-rod dystrophy, or even an isolated cone dystrophy (Grudzinska Pechhacker et al., 2021). Patients with BBS10 may present with simultaneous night blindness and peripheral field constriction, as well as reduced central vision, a devastating combination (Grudzinska Pechhacker et al., 2021). *Bbs10^-/-^* mice demonstrate a similar phenotype making them an excellent avatar for preclinical treatment trials. It is worth noting that the mouse models used in this paper are not both knockouts. The BBS10 mouse model is a full knockout that produces no *Bbs10* mRNA while the BBS1 mouse model is a substitution mutation changing just one amino acid in the protein. This may contribute to the difference in severity seen between the mouse models. However, this is an accurate comparison of the two diseases for humans. The most common cause of BBS10 in humans is a single nucleotide insertion that results in premature termination and loss of protein expression (Stoetzel et al., 2006), while the most common cause of BBS1 is this same amino acid substation mutation reported in the mouse model here (Davis et al., 2007; Mykytyn et al., 2002). This substitution mutation may leave some part of a functioning protein, albeit abnormal while the BBS10 mutation lacks any protein whatsoever. Thus, a partially functional *BBS1* gene may result in a partially functional BBSome, while a completely nonfunctional *BBS10* gene may lead to little or no BBSome formation, explaining the more severe BBS10 retinal phenotype in mouse and man.

In conclusion, a knockout mouse model of BBS10 recapitulates the human retinal degeneration and shares features of the human condition. Comparison to a model of BBS1 demonstrates that different subtypes of BBS will likely require different rescue strategies and timing of treatment. Rods and cones are present early in life in *Bbs10^-/-^* mice retinas but are abnormal and progressively degenerate over time. The early non-recordable 5 Hz flicker ERG offers a robust endpoint for therapeutic rescue studies. Maintaining useful functional vision in the light even when ERGs are nonrecordable suggests that rescuing even a small percentage of cones could provide useful visual improvement to patients.

## Materials and Methods

### Animals

Experiments were approved by the Institutional Animal Care and Use Committee (IACUC) at the University of Iowa and conducted following the recommendations in the Guide for the Care and Use of Laboratory Animals of the National Institutes of Health. Mice were maintained on a standard 12/12 hour light/dark cycle with food and water provided *ad libitum.* The breeders to start the line contained the *Rd8* mutation in *Crb1;* this was bred out early in the process. Genetic testing confirmed WT for the *Crb1* gene as well as genotyping for the knockout mutation. Wild type 129/SvJ mice were obtained from Jackson Laboratory, Bar Harbor, Maine.

### Optical Coherence Tomography

OCTs were performed using the Bioptigen OCT with a small rodent lens (Research Triangle Park, NC) as previously described(Drack et al., 2012). Mice were anesthetized with a mixture of ketamine (87.5 mg/kg) and xylazine (2.5 mg/kg) and pupils were dilated with tropicamide 1%. Measurements were taken of the outer nuclear layer as measured from 3μm on either side of the optic nerve on all images. The measurements were done using the in-software calipers provided by Bioptigen.

### Microscopic anatomy

Mice were sacrificed at times indicated. Retinal sections were stained with rabbit anti-OPN1MW polyclonal antibody (EMD Millipore AB5405) and rabbit anti-GNAT2 polyclonal antibody (Abcam ab97501). Methods to perform immunofluorescence stainings are as previously described (Datta et al., 2021; Datta et al., 2019; Datta et al., 2020). Transmission electron microscopy was performed as described previously (Hsu et al., 2017).

### Electroretinograms

Full-field electroretinograms (ERGs) were conducted within 2 weeks of each swim assay. Full-field ERG was performed using the Celeris Diagnosys system (Diagnosys LLC, Lowell, MA). The mice were dark adapted overnight before ERG was conducted. Mice were anesthetized with a mixture of ketamine (87.5 mg/kg) and xylazine (2.5 mg/kg). The mice received 0.1 mL of the mixture per 20 g body weight. ERGs were recorded simultaneously from the corneal surface of each eye after the pupils were dilated with 1% tropicamide, using Diagnosys Celeris touch stimulator electrodes. Gonak gel (Akorn, Inc., Lake Forest, IL) was place on the cornea of each eye before the electrode was positioned. Light flashes were produces by the touch stimulator electrodes. Dim red light was used for room illumination until dark-adapted testing was completed. A modified International Society for Clinical Electrophysiology of Vision (ISCEV) protocol was used. A dim flash measured at 0.01 cd•s/m^2^ is performed first to measure rods, followed by a bright flash at 3.0 cd s m-2 to measure the standard combined response (SCR) of the rods and cones. Mice are then light adapted for 10 minutes, after which they are tested with a bright flash at 3.0 cd•s/m^2^ followed by a flicker of 5 Hz. 15 sweeps are taken of each condition except the 5Hz flicker, in which 20 sweeps are taken. ERG results are analyzed following data collection using the Diagnosys software to eliminate sweeps that show interference such as mouse movements or ambient electric signal. If more than 5 sweeps needed to be eliminated, the test was null and was repeated for cleaner results. The light adapted protocol tests cones, especially in the case of the 5 Hz flicker which only elicits a response from cones in mice(Tanimoto et al., 2015).

### Visually Guided Swim Assay

A modified Morris water maze (Morris, 1984) incorporating some features of the mouse swim assay developed by Pang et al. (Pang et al., 2008) was utilized to test functional vision. A plastic swimming pool was used for swimming. The pool diameter measures 35 inches at the bottom, 38 inches at the 4-inch water level, and 39 inches at the top. A 3” diameter pvc tube with rubber caps on both ends was used as a platform for the mice to climb onto to end the trial. The platform was the same on both ends, so it could be flipped between trials to mitigate any odor that might aid in platform localization. A “flag”, consisting of a small paper American flag taped to a wooden stick was attached to the side of the platform to make it more visible to the mice. Platform location consisted of 8 equally spaced pre-set locations. The order of the locations of the platforms was determined prior to the day’s experiment and was randomly determined each day.

Swim protocol consists of 4 light adapted training days followed by 4 light adapted testing days. Following light adapted testing, 2 days of dark adapted training and then 4 days of dark-adapted testing are performed. Light adaptation was performed in a brightly lit room of typical fluorescent ceiling lights. Dark adaptation was performed in the same room with the lights off and dim red lights to assist the observers to see the experiments. An infrared lamp was aimed at the pool and the location of the mice was determined by night vision goggles.

Each testing day of swimming, mice performed 5 swim trials to 5 different randomly selected platform locations. These platforms were the same for all mice in the testing group. A limit of 60 seconds was set for swim TTPs to prevent mice from becoming fatigued; if the platform was not attained by 60 seconds, the mouse was gently removed from the pool, and the time to platform for that particular trial was recorded to be 60 seconds.

### Genotyping

Genotyping was performed using mouse tail snips collected at P14. DNA was extracted and *Bbs10* was amplified using wildtype and knockout forward primers and the same reverse primer (Supplementary materials).

### RT-PCR

RNA was extracted from whole eye lysates and amplified up using SuperScript VILO Master Mix to create a cDNA library. After this, PCR was run on the cDNA amplified by primers indicated (Supplementary materials). Actin was used as a control.

### Statistical Analysis

Statistical analysis and figure creation was done using GraphPad Prism. Comparisons between age group means was done using a Sidak’s multiple comparison’s test for ERG, OCT and weight data. *Bbs10^-/-^* mice were compared to littermate controls for all experiments. Graphs for weight data and OCT were plotted using mean and standard deviation. Graphs for ERG was plotted using the mean and standard error of the mean (SEM). Comparison p-values between TTP in the swim assay were calculated using a Student’s T-test. All trial swim times were used to calculate the p-values and figures were made using the mean and SEM. All analyses were performed using an alpha value of 0.05.

## Acknowledgements

We thank L.L. Wallrath for edits to the manuscript. This research was supported by grants from Fighting Blindness Canada, The Bardet Biedl Syndrome Family Association, Ronald Keech Professorship, and University of Iowa Institute for Vision Research. This research was supported in part by an NIH/NEI Center Support Grant to the University of Iowa (P30 EY025580).

## Author contributions

Manuscript writing and figure preparation: S.K.M., J.T., Y.H. and A.V.D.

Data Acquisition: S.K.M., J.T., M.H., A.K., I.C., A.S., E.S., P.D., C.S., Y.H., S.B. Materials provided: C.S., S.S., V.C.S.

Study concept and design: S.S., V.C.S., A.V.D.

Statistical analysis: S.K.M., K.W.

Additional Information/Competing Interests: NA

---

**Figure S1:**
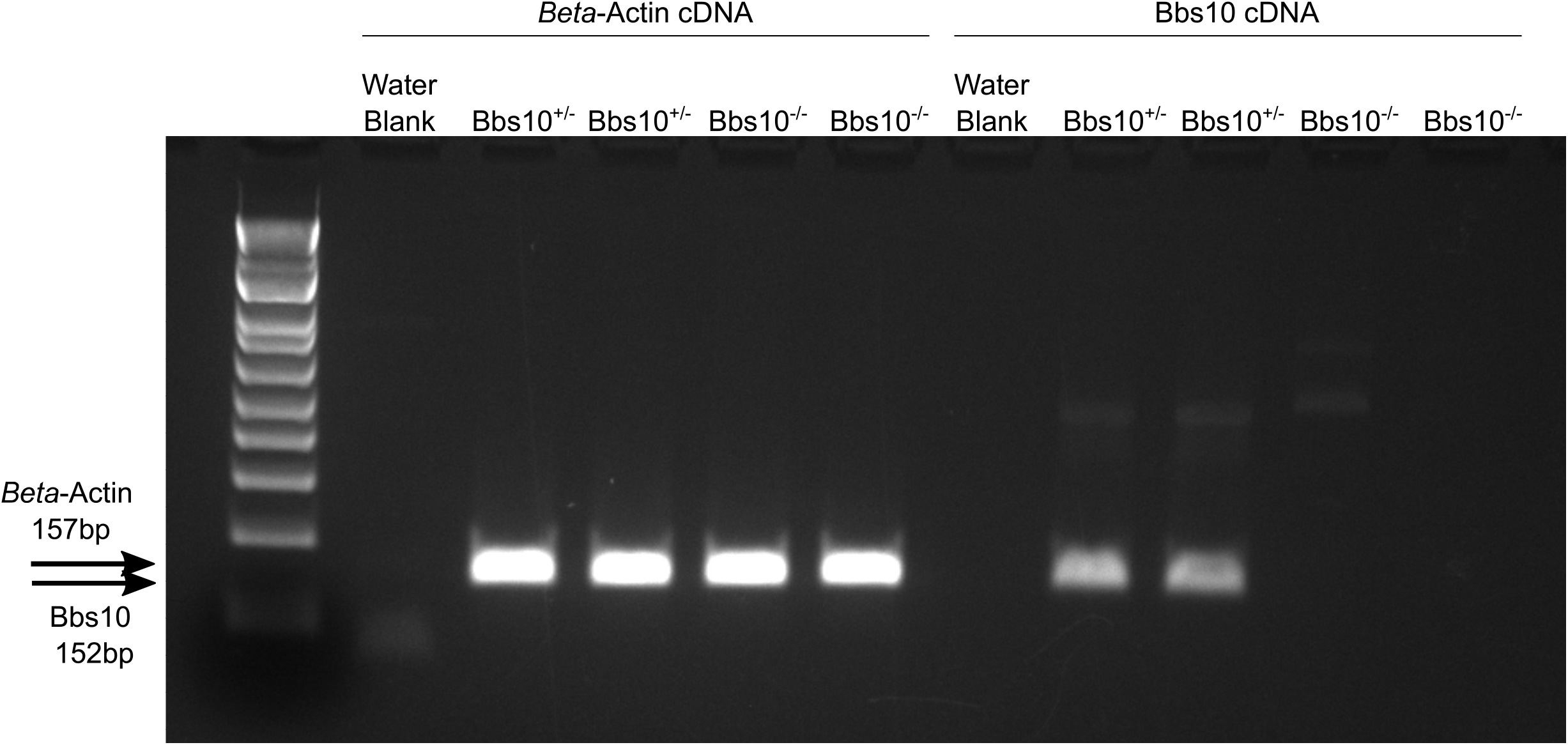
Bbs10^-/-^ mice have no Bbs10 mRNA present. The figure displays RT-PCR results demonstrating that Bbs10^-/-^ do not produce any mRNA that can be detected. Actin was used as a control, which shows robust expression data in all mice.

**Figure S2:**
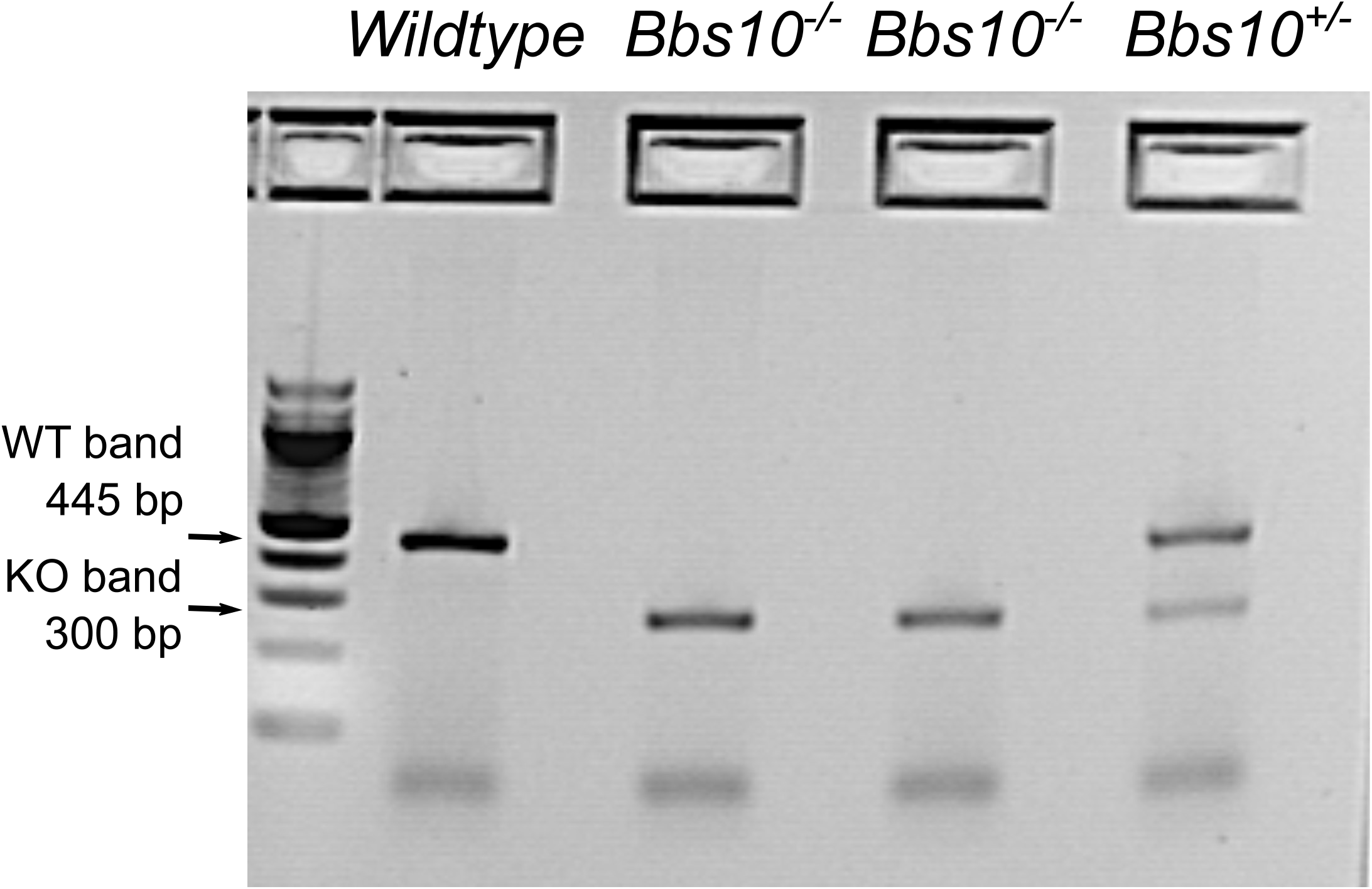
*Bbs10^-/-^* mice required genotyping due to breeding limitations of *Bbs10^-/-^* males. *Bbs10^-/-^* mice have a deletion of both exons of the *Bbs10* gene and thus have a smaller PCR product upon genotyping that created bands located at 445bp for the WT band and 260bp for the knockout band. Two appropriately sized products are present in heterozygous mice.

**Figure S3:**
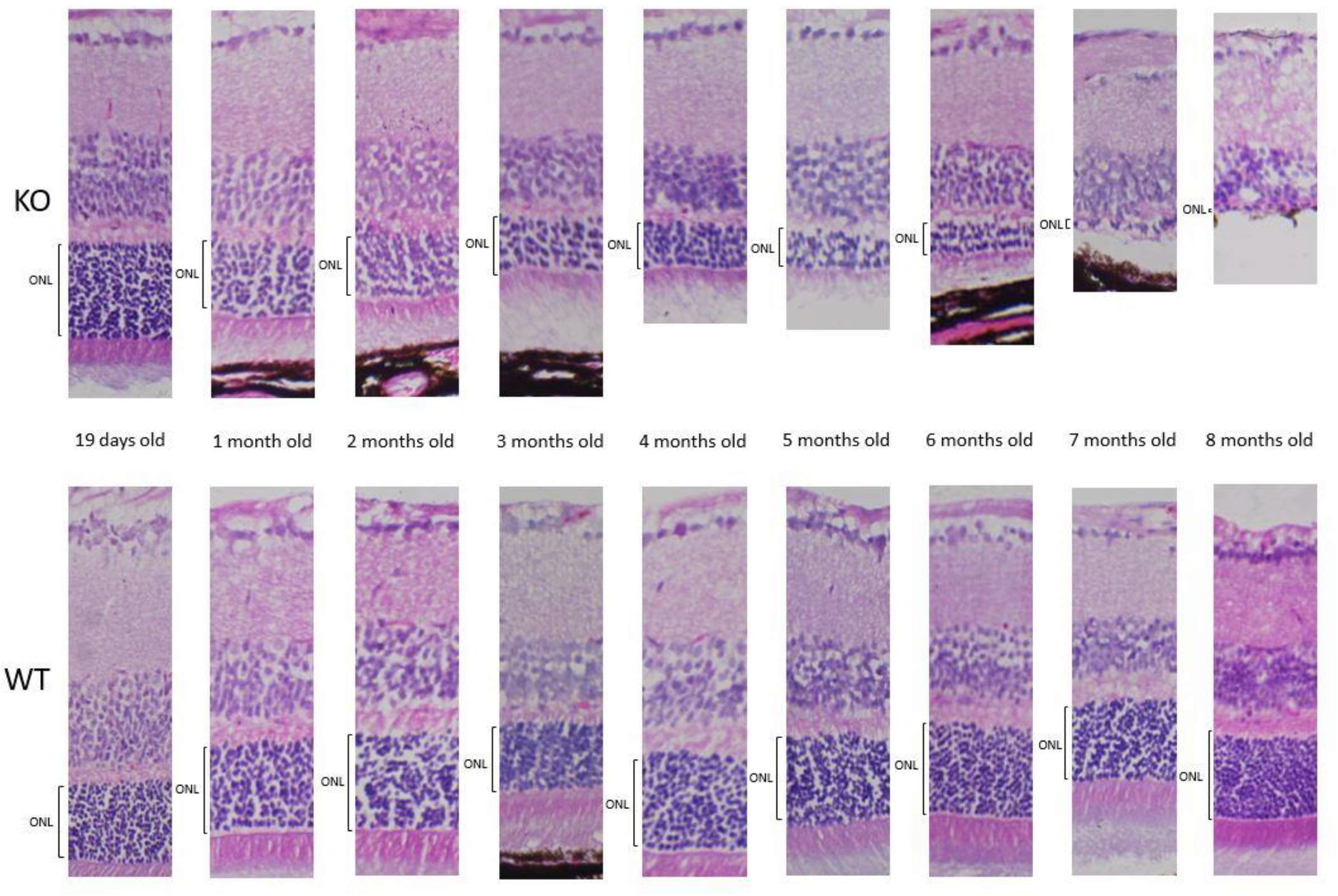
*Bbs10^-/-^* mice have a thinner outer nuclear layer (ONL) compared to control mice. Control mice have a constant thickness of the ONL throughout their lifespan. Differences in apparent thickness in control images is due to sectioning techniques that spreads out the cell layers. In contrasts, the ONL layer of *Bbs10^-/-^* mice become progressively thinner over the mouse lifespan and show only a single-cell layer by seven months of age.

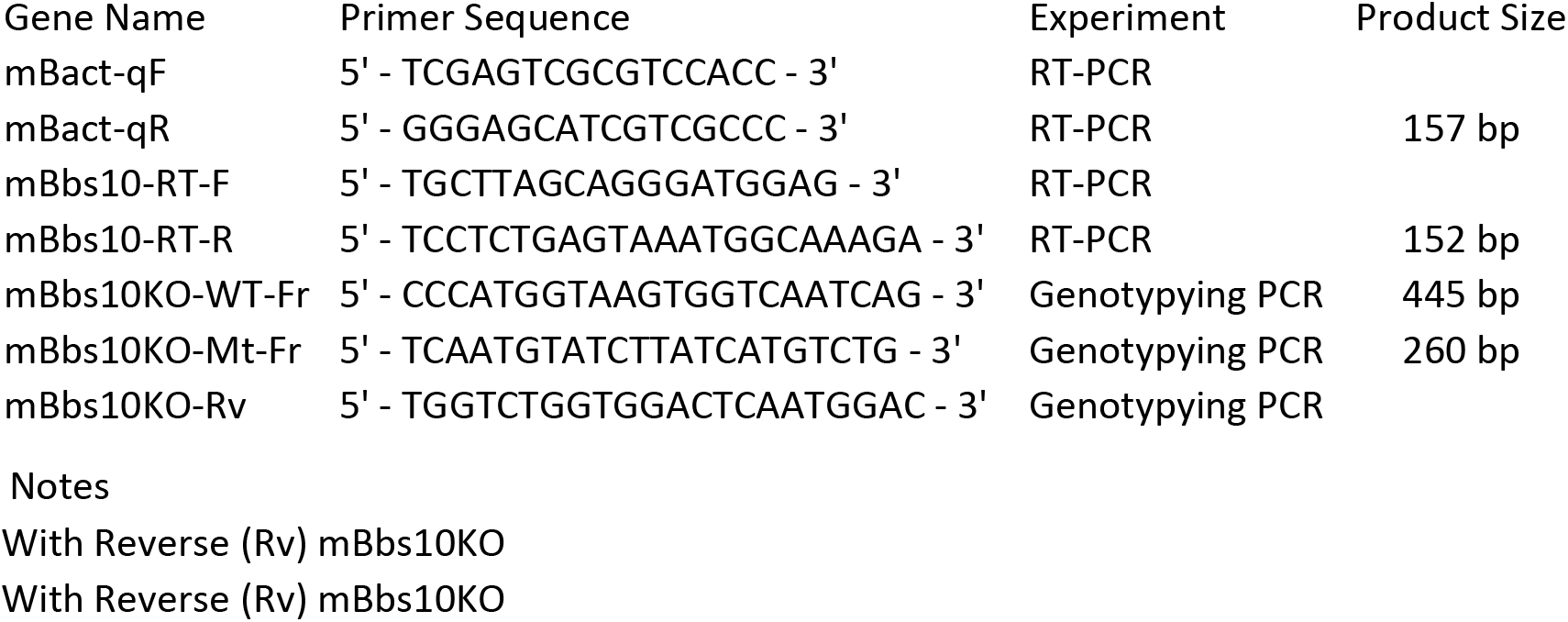

